# Screening of epigenetic modifiers identifies novel host-directed agents that suppress HIV-1 replication in primary human macrophages

**DOI:** 10.64898/2026.09.01.748500

**Authors:** Gabrielle Lê-Bury, Christine Mandalasi, Jordan M. Rhen, David W. Gludish, Saikat Boliar, Henry C. Mwandumba, David G. Russell

## Abstract

Epigenetic modifications play a critical role in diverse biological processes, including HIV- 1 replication in lymphoid and myeloid cells, particularly in the establishment and maintenance of latency. As such, epigenetic targets represent potential candidates for novel host directed therapy (HDT). In this study, we developed a novel *in vitro* screening platform using human primary macrophages, including both monocyte-derived macrophages and alveolar macrophages, infected with a replication-competent luminescent virus. In contrast to previous screens, this platform enables the identification of compounds and epigenetic targets that modulate viral replication during a spreading infection, capturing not only transcriptional regulation but other vulnerabilities across the complete HIV-1 life cycle. Our data reveal multiple novel compounds targeting distinct epigenetic regulators, effectively inhibiting viral replication at various stages in primary macrophages, which may serve as potential candidates/targets for HDT.

## INTRODUCTION

After more than 40 years of research, HIV-1 (Human Immunodeficiency virus type-1) remains a major global public health challenge, with more than 40 million people living with the virus in 2024 (WHO, 2025). In 1987, Azidothymidine (AZT), a nucleoside-analog reverse transcriptase inhibitor (NRTIs), became the first FDA-approved antiretroviral drug to treat HIV/AIDS (Fauci and Lane 2020). Since then, numerous antiretroviral drugs have been approved by the FDA, distributed across several classes: (i) NRTIs; (ii) nonnucleoside reverse transcriptase inhibitors (NNRTIs); (iii) integrase inhibitors; (iv) protease inhibitors; (v) fusion inhibitors; (vi) coreceptor antagonists; and more recently (vii) capsid inhibitor (Fauci and Lane 2020; Segal-Maurer et al. 2022). The actual regimen for combination antiretroviral therapy (cART) is mostly based on direct-acting antivirals (DAA) that suppress HIV-1 replication to an undetectable level and prolong the lifespan of infected individuals (Pagani, Ghezzi, et al. 2022). However, the emergence of drug-resistant viruses poses a challenge to the current cART regimens which could be further bolstered with additional classes of antiretrovirals. One novel approach for antiviral drug development is host-directed therapy (HDT) emphasizing drugs modifying epigenetic regulation of the host genome (Kumar et al. 2020; Shepherd et al. 2025).

Epigenetic modifications play a crucial role in HIV-1 persistence within various viral reservoirs including myeloid cells (Verdikt, Hernalsteens, and Van Lint 2021; Krause, Bergmann, and Schmidt 2024). These epigenetic mechanisms, including DNA methylation, histone modifications and non-coding RNAs, are known to regulate HIV-1 latency (Boliar and Russell 2021; Nguyen and Karn 2024). One proposed therapeutic approach is the “shock and kill” strategy using latency-reversing agents (LRAs) to reactivate HIV-1 gene expression in lymphoid and myeloid viral reservoir cells (Ait-Ammar et al. 2019). For instance, non-coding RNAs known to be involved in HIV-1 latency, might be genetically or chemically targeted to control HIV-1 persistence (Boliar and Russell 2021; Le-Bury et al. 2023; Beliakova-Bethell 2024). Macrophages are more resistant to the cytopathic effects of HIV-1 infection than T cells. Several cellular factors including long non-coding RNAs (lncRNAs) are known to induce a pro-survival response in HIV-1-infected cells. It has been shown that and treatment with siRNAs targeting the lncRNA SAF drives cell death in infected but not in uninfected macrophages (Boliar et al. 2019). This demonstrates that targeting cellular pathways represents a potential new avenue of viral control.

There is a growing appreciation of the significance of tissue resident macrophages as sites of viral persistence in HIV-1 infection (Ganor et al. 2019; Hendricks et al. 2021; Pagani, Demela, et al. 2022). Epigenetic regulation differs between HIV-1-infected T cells and macrophages, suggesting that these cell types need to be studied independently (Deshiere et al. 2017; Lu et al. 2022). A recent study utilized human monocyte-derived macrophages (hMDMs) to perform a high throughput screen with a collection of 6,000 small molecules and identified compounds that can inhibit viral transcription (Yi et al. 2023). The infection was initiated with a Δenv, luciferase- expressing virus that was pseudotyped with VSVg-glycoprotein that generates a single round of infection and therefore limited to a non-spreading infection model. Nonetheless, the screen identified a purine nucleoside analog, nelarabine as an inhibitor of viral transcription that also reduced the levels of H3K9meK marks on the provirus indicating that the nucleoside analog might impact viral fitness through epigenetic modulation (Yi et al. 2023).

In this current study, we interrogated the impact of epigenetic modulation on a replication-competent, spreading HIV infection. We performed a high throughput screen to identify epigenetic modulators that impacted HIV-1 infection in primary hMDMs and lung-resident alveolar macrophages. There are several features in the design of this study that are of particular significance. First, we used intact wild-type virus with relatively low multiplicity of infection (MOI) to initiate infection, thereby establishing a spreading viral infection that recapitulates the complete viral life cycle. Viral replication was measured using bioluminescence from viral LTR-driven expression of Gaussia luciferase and is referred to as “viral productivity”. This contrasts with the majority of similar chemical screening assays conducted on lymphocytes and macrophages, where a single round of infection is commonly interrogated. Secondly, we utilized a focused library of 721 compounds from MedChemExpress (MCE) as a collection of epigenetic inhibitors/activators that have putative known targets relevant to cellular re-programming pathways. From the screen we identified several classes of epigenetic modulators and inhibitors that reduced viral productivity. These include both known anti-HIV-1 compounds such as JQ-1 as well as previously unknown inhibitors such as dBET1. This study has identified several points of vulnerability within the viral replication cycle that may represent tractable targets against HIV-1 for future drug development.

## RESULTS

### Macrophage-based screening identifies modulators of HIV-1 replication in primary cells

To identify pathways of epigenetic regulation within the HIV-1 replication cycle in human macrophages, we developed a screening platform using a library that was enriched for epigenetic modifiers acquired from MCE. This library includes 721 small molecules across 16 different classes of epigenetic modifiers: epigenetic reader domain (BRD); histone methyltransferases (HMT); histone deacetylases (HDAC); Janus kinases (JAK); poly(ADP-ribose) polymerases (PARP); protein kinases (PKC); HIF/HIF proxyl-hydroxylase (HIF); AMP-activated protein kinases (AMPK); Sirtuins (SIRT); Aurora kinases (AURK); histone demethylases (HDM); histone acetyltransferases (HAT); DNA methyltransferase (DNMT) ; Pim kinases (Pim); protein arginine deiminase (PAD); and microRNA (Fig 1A). We engineered a virus with NL4-3 backbone and R5- tropic Env (BaL) to express Gaussia luciferase (NL43-BaL-IRES-GLuc) to evaluate the effect of the epigenetic compounds on a productive and spreading infection in primary macrophages (Fig 1B). The readout of viral replication was total luciferase activity, which is effectively a composite output of all the biological processes in the viral replication cycle, and as such is referred to as “viral productivity”. In hMDMs, the integration of the viral genome occurs up to 4 days post-infection (dpi) (Yi et al. 2023; Bejarano et al. 2018; Castellano et al. 2019). Thus, cell treatment one day post infection allows us to evaluate the effect of compounds on different steps of viral cycle (reverse transcription, integration and viral transcription, viral budding and spreading infection), with minimal manipulation.

**Figure 1:**
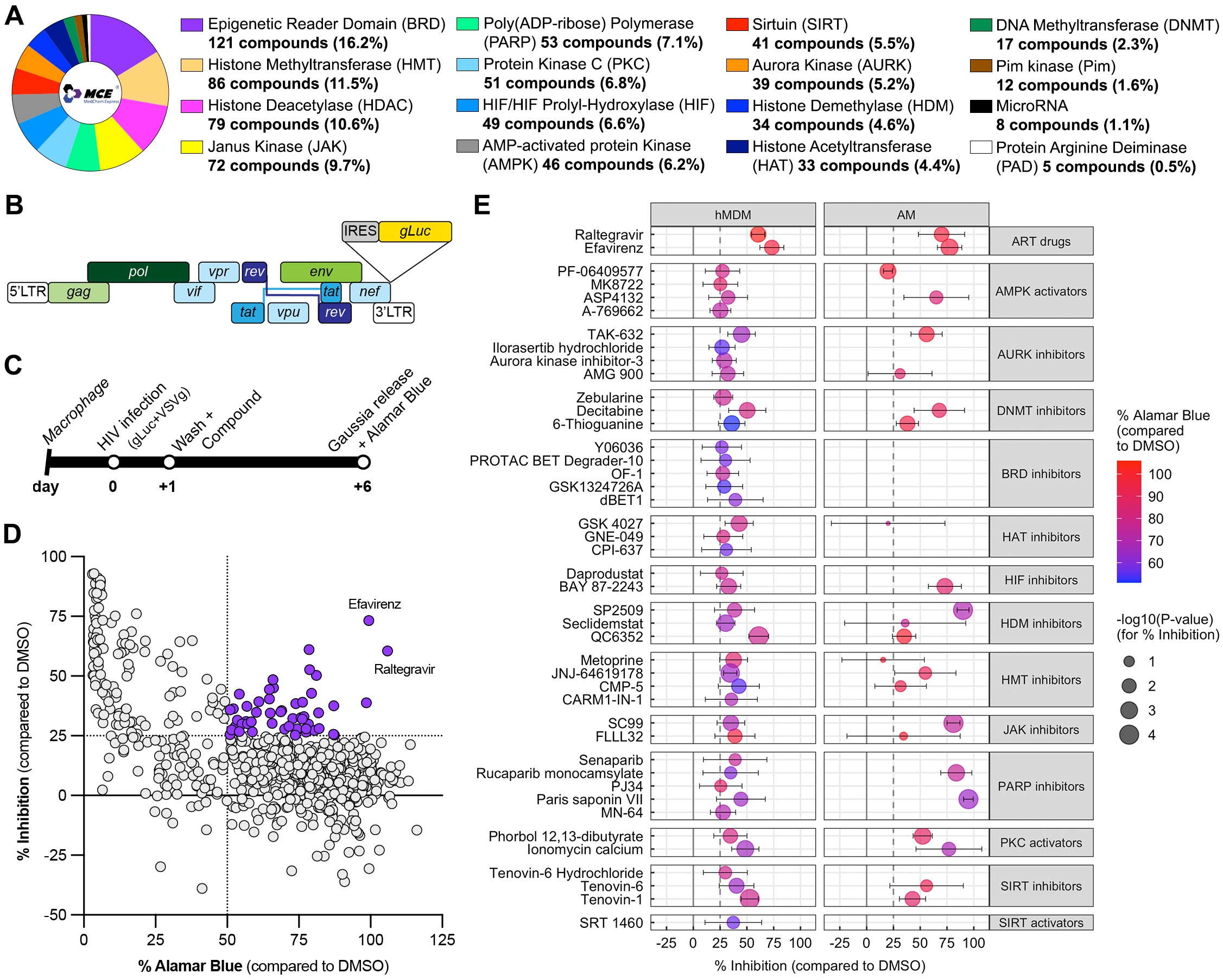
Identification of epigenetic modulators of HIV replication in primary macrophages. (A) Composition of the Epigenetic Small Molecule Library from MedChemExpress encompassing 721 compounds from 16 different target classes. (B) Schematic of the HIV-1 reporter construct NL4-3-IRES-gLuc with R5-tropic env (BaL). (C) Schematic of experimental design of the macrophage-based screening assay in primary cells. Day 0, macrophages were infected with HIV-gLuc pseudotyped with VSVg at MOI of 0.1, then washed and treated with epigenetic compounds at 10mM the following day. After 5 days of treatment, supernatants were collected for Gaussia luciferase quantification and Alamar Blue was used to evaluate viability. (D) Graphical representation of screening data with the percentage of inhibition of viral replication (% Inhibition) and the percentage of viability (% Alamar Blue) compared to HIV-infected cells treated with DMSO, in y-axis and x-axis, respectively. Each dot represents the average of each compound from 5 different donors, with compounds showing more than 25% inhibition and 50% viability in purple. (E) Graphical representation of % Inhibition (x-axis), % Alamar Blue (color scale) and the *p*-value of % Inhibition (dot size) for 41 selected compounds (purple in D) with their names (far left) and their epigenetic classes (far right) in human monocyte-derived macrophages from 5 donors (left) and alveolar macrophages from 4 donors (right).

Following 7 days of differentiation, hMDMs were detached; seeded in 96-well plates; infected the next day with HIV-Gluc pseudotyped with VSVg (Vesicular Stomatitis Virus glycoprotein) at MOI 0.1; washed 1dpi, and test compounds added at 10mM, which included Efavirenz (EFV, NNRTI) and Raltegravir (RAL, integrase inhibitor) as positive controls. Viral productivity and host cell fitness were assessed at 6dpi by measuring Gaussia luciferase release in the supernatant and Alamar Blue quantification, respectively (Fig 1C). We identified 41 small molecules that significantly inhibited bioluminescence signal by over 25%, while maintaining more than 50% of the Alamar Blue signal (Fig 1D), as a measure of cellular viability. To ensure we incorporated potential host macrophage heterogeneity between monocyte-derived (hMDMs) and tissue resident lineages, we also assessed the inhibition efficacy of several of the 41 hits on viral productivity in alveolar macrophages (AMs) acquired through bronchoalveolar lavage (BAL) from 4 healthy donors. After isolation, AMs were plated directly in 96-well plates, infected 4h later with VSVg pseudotyped HIV-Gluc *ex vivo* and treated with compounds 1dpi, consistent with the hMDM infection (Fig 1C). In AMs, the variability was higher, however, most of the hits identified in the primary screen on HMDMs inhibited viral productivity greater than 25% (Fig 1E). Interestingly, some compounds suppressed the luciferase signal in AMs more effectively than in hMDMs (Fig 1E). Thus, the hMDM-based screen identified a diverse collection of epigenetic regulators that restricted HIV productivity in primary human macrophages, both monocyte-derived and tissue resident (Table S1). To begin to probe the mode of action (MOA) of these compounds, we designed several secondary screens to assess the stages of the HIV-1 life cycle that were impacted by the inhibitors.

### Screening of infected primary macrophages treated after viral genome integration identifies modulators of HIV-1 transcription

HIV-1 infection of cells has a major impact on chromosomal accessibility and transcriptional activity, and epigenetic modifications are among the most important regulators of viral transcription (Verdikt, Hernalsteens, and Van Lint 2021). To assess the efficacy of selected compounds on viral transcription in macrophages, we infected hMDMs with HIV-Gluc and initiated treatment with epigenetic compounds at 4 dpi, after integration had occurred (Supp Fig 1). Medium was supplemented with the CCR5 antagonist Maraviroc (MVC) to prevent reinfection. Two days later, Gaussia luciferase activity and Alamar Blue assays were performed to assess viral transcription and host cell fitness, respectively (Fig 2A). We identified two compounds that increased viral transcription by more than 25%, similar to the latency-reversing agent (LRA), Bryostatin (PKC agonist), which was included as a positive control: these were Phorbol 12,13- dibutyrate (PDBu), another PKC activator; and Seclidemstat, a HDM inhibitor targeting LSD1 (lysine-specific histone demethylase 1) (Fig 2B). Conversely, two compounds reduced viral transcription by over 25%, to a level comparable to the known transcription inhibitor, Flavopiridol: these were Senaparib, a PARP inhibitor, and dBET1, a BRD4 inhibitor (Fig 2B). Cells treated with Senaparib and dBET1 significantly reduced viral transcription to 34% and 38%, respectively (Table S1). Amongst the PARP and BRD4 inhibitors of the MCE library, Senaparib and dBET1 were the most efficient inhibitors of viral productivity with these cellular targets (Table S1).

**Figure 2:**
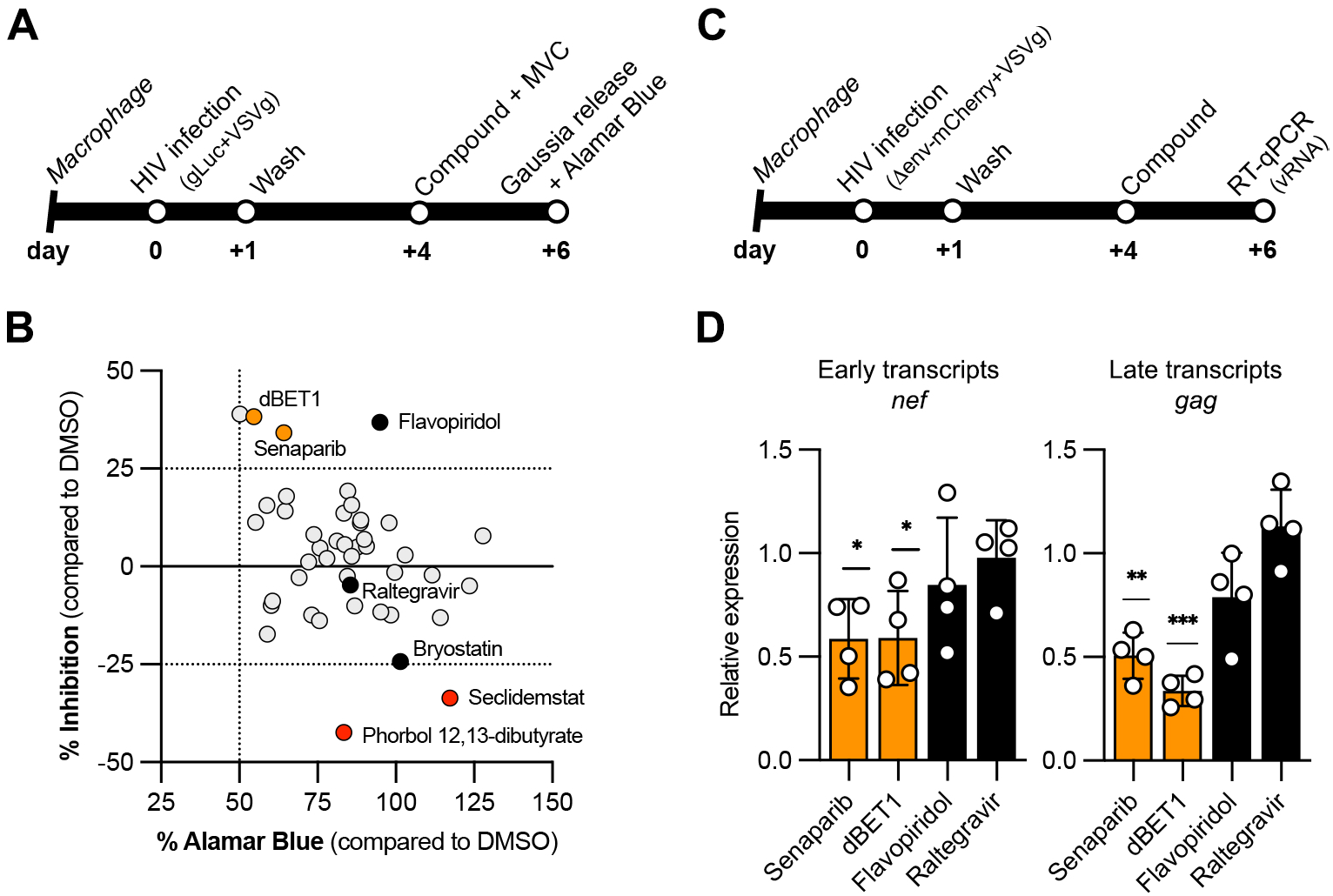
Identification of epigenetic modulators controlling HIV transcription in hMDMs. (A) Schematic of experimental design of the silencing screening assay in hMDMs. Macrophages were infected with HIV-gLuc pseudotyped with VSVg at MOI of 0.1, then washed the following day. After 4 days of infection, cells were treated with Maraviroc (2mM) and epigenetic compounds (10mM), then supernatants were collected 2 days later for Gaussia luciferase and Alamar Blue quantification. (B) Graphical representation of the data with the percentage of inhibition of viral replication (% Inhibition) and the percentage of viability (% Alamar Blue) compared to HIV-infected cells treated with DMSO, in y-axis and x-axis, respectively. Each dot represents the average of a compound from 3 different donors, with in orange, compounds showing more than 25% inhibition and 50% viability, and in black, controls (Flavopiridol, Raltegravir, Bryostatin). (C) Schematic of experimental design of the silencing assay in primary cells. Macrophages were infected with non-replicative HIV-NL4-3-Denv-mCherry pseudotyped with VSVg at MOI of 0.2, then washed the following day. After 4 days post-infection, cells were treated with Senaparib or dBET1 at 10mM, or controls (Flavopiridol at 100nM or Raltegravir at 10mM). Two days later, cells were lysed for RNA extraction and RT-qPCR. (D) RT-qPCR analysis of *nef* (left) and *gag* (right) mRNA for Senaparib and dBET1 (orange bars), and controls (black bars) compared to HIV-infected cells treated with DMSO (grey bar) from 4 donors.

To confirm the that the effect of these two compounds was on viral transcription, we used a *env*-negative virus pseudotyped with VSVg, treated the cells at 4 dpi and extracted RNA at 6 dpi (Fig 2C). The RT-qPCR data showed that Senaparib inhibited the viral transcription of *nef* and *gag* genes at 58.64% and 50.63%, respectively, while dBET1 reduced *nef* and *gag* gene expression at 59.03% and 33.64%, respectively (Fig 2D). These results indicate that the primary screen (Fig 1) identified compounds that reduce viral productivity though a MOA that impacts viral transcription.

### Screening of primary macrophages treated prior to viral infection identifies inhibitors of HIV-1 reverse transcription

Another stage in the viral life cycle that is well known as being vulnerable to chemical inhibition is reverse transcription (RT) (Singh and Das 2022). To evaluate the effect of compounds on the earlier steps of viral replication in human primary macrophages, cells were treated one day prior to infection, then washed, infected, and treat recommenced one day post-infection (Fig 3A). From the 41 active compounds, 28 compounds exhibited minimal impact on hMDM fitness (Alamar Blue>50%) including 22 compounds that inhibited viral productivity by more than 50% (Fig 3B). These compounds included several different classes of epigenetic modifiers, and most of those tested also inhibited viral productivity in HIV-infected AMs when the AMs were treated similarly, before *ex vivo* infection (Fig 3C). To assess the relative potency of these compounds, we generated dose response curves for the 22 selected small molecules and recorded both IC_50_ values and Alamar Blue signal (Supp Fig 2). Consistent with the initial screen, these compounds were more efficient if macrophages were treated one day before infection as opposed to one day post infection (Fig 3D).

**Figure 3:**
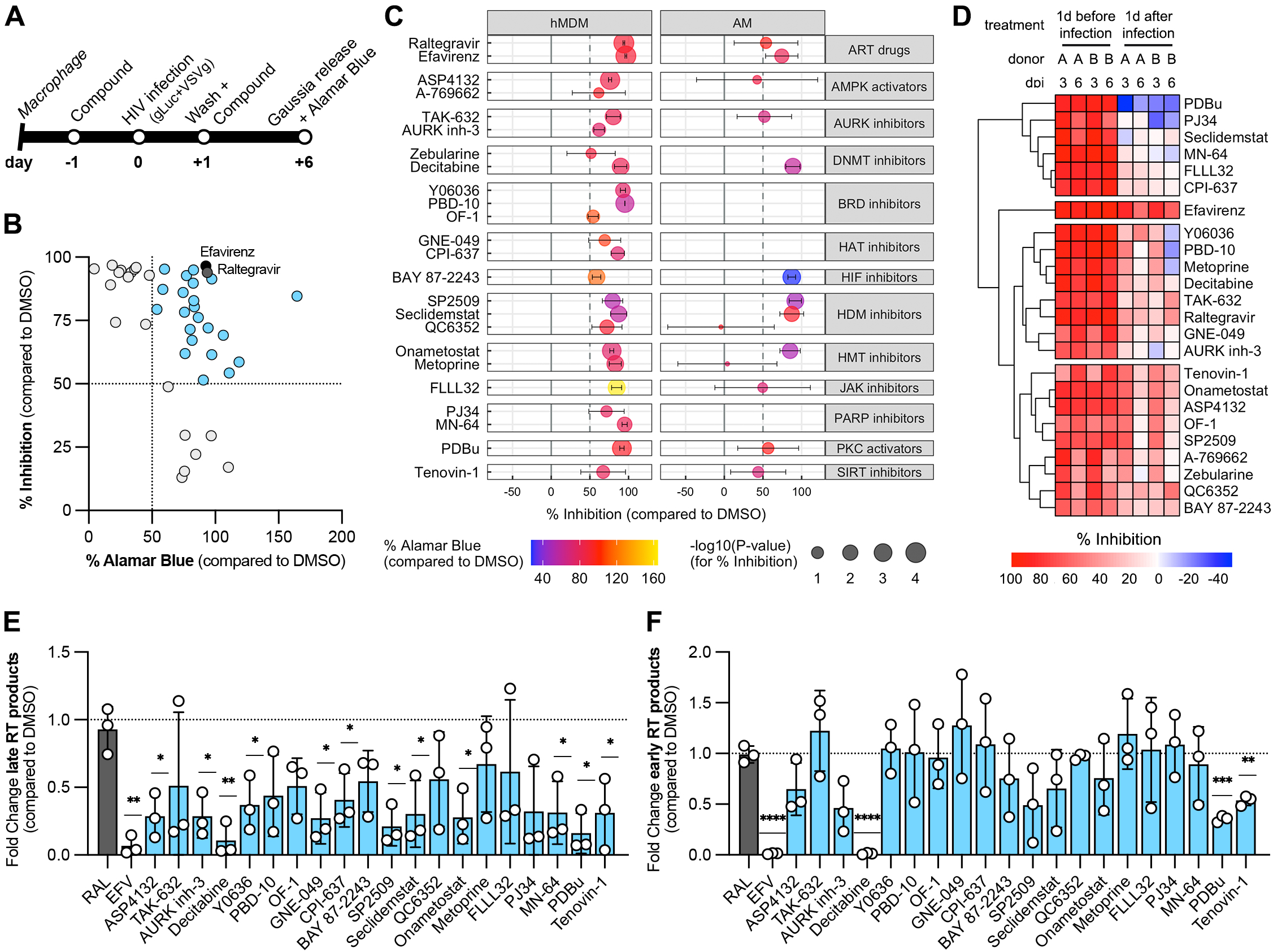
Identification of epigenetic compounds regulating reverse transcription in hMDMs. (A) Schematic of experimental design of the screening assay in primary cells. Macrophages were treated with epigenetic compounds at 10mM, then infected with HIV-gLuc pseudotyped with VSVg at MOI of 0.1 the next day. The following day, cells were washed and treated with epigenetic compounds, then, 6 days post-infection, supernatants were collected for Gaussia luciferase and Alamar Blue quantification. (B) Graphical representation of the data with the percentage of inhibition of viral replication (% Inhibition) and the percentage of viability (% Alamar Blue) compared to HIV-infected cells treated with DMSO, in y-axis and x-axis, respectively. Each dot represents the average of one compound from 3 different donors, with in light blue, compounds showing more than 50% inhibition and 50% viability, and in black and dark grey, the positive controls Efavirenz and Raltegravir, respectively. (C) Graphical representation of % Inhibition (x-axis), % Alamar Blue (color scale) and the *p*-value of % Inhibition (dot size) for the 22 selected compounds (light blue in B) with their names (far left) and their epigenetic classes (far right) in human monocyte-derived macrophages from 3 donors (left) and alveolar macrophages from 3 donors (right). (D) Macrophages were treated with the 22 selected epigenetic compounds at 10mM before (left) or after (right) HIV-gLuc infection (MOI 0.1), and % Inhibition (color scale) was calculated at 3dpi and 6dpi for 2 donors (donor A and donor B). (E-F) Macrophages were treated one day before HIV-gLuc infection (MOI 0.2) and 2dpi DNA was extracted to quantify late (E) and early (F) RT products compared to HIV-infected cells treated with DMSO, for 3 donors.

To determine whether any of these inhibitors impacted RT, cells were infected with a non-replicative virus (NL4-3 Denv mCherry pseudotyped with VSVg), then treated with compound the next day (Fig. 3A), and viral DNA was measured by qPCR at 2dpi. The reverse transcription process is characterized by different stages − initiation, minus-strand transfer, elongation and completion − which can be quantified by qPCR using different pairs of primers (Mbisa et al. 2009). In addition, it has been reported previously that RT activity in macrophages is slower and the stages more distinct than reported for other cell types (Collin and Gordon 1994). For this reason, we performed qPCR on early and late RT intermediates as detailed (Mbisa et al. 2009). Half of the compounds (12 out of 22) significantly inhibited production of late RT products by more than 50% (Fig 3E). However, only 3 compounds significantly reduced early products of RT by more than 45%: Decitabine (DNMT inhibitor), Phorbol 12,13-dibutyrate (PKC activator), and Tenovin-1 (SIRT inhibitor) (Fig 3F). The data confirm that the macrophage-based screen (Fig 1C-D, Fig 3A- B) successfully identified several new compounds impacting reverse transcription. It is notable that amongst these compounds is an inhibitor known to impact multiple cellular functions, Tenovin-1 (Ladds et al. 2020; Lain et al. 2008).

### Tenovins-1 inhibits viral replication at RT and post-RT steps in primary macrophages

Tenovin-1 has been reported to inhibit both Zika and Dengue virus replication (Rausch et al. 2017; Hackett et al. 2019; Wan et al. 2021). In our study, screens performed on hMDMs and AMs, also detected inhibitory activity on viral replication mediated by both Tenovin-1 and its analogs, Tenovin-6 and -6 hydrochloride, (Fig 1E). In AMs, treatment with Tenovins pre- or post-infection showed minimal cytotoxicity (64% to 94% Viability), and Tenovin-6 appeared to be more potent than Tenovin-1, with 49% to 58% and 38% to 55% of inhibition of viral productivity, respectively (Fig 4A). In contrast, in hMDMs, Tenovin-1 was more potent than Tenovin-6 and Tenovin-6 hydrochloride with treatment post-infection, and also exhibited lower cytotoxicity following treatment pre-infection (Fig 4A). To confirm these observations, we performed dose responses experiments by pre-treating hMDMs with Tenovins prior to HIV-gLuc infection (Fig 3A). At 6dpi, Tenovin-6 and Tenovin-6 hydrochloride showed an IC_50_ at 1.3mM and 2.6mM, respectively, while IC_50_ of Tenovin-1 is at 4.9mM (Fig 4B, Table S1). The CC_50_ of Tenovin-1 could not be determined because of its minimum cytotoxicity, while Tenovin-6 and Tenovin-6 hydrochloride showed a CC_50_ at 23.3mM and 28mM, respectively (Fig 4B, Table S1). These results indicate that Tenovin-1 is more efficient than Tenovin-6 and Tenovin-6 hydrochloride at inhibition of viral replication, and shows minimal cytotoxicity.

**Figure 4:**
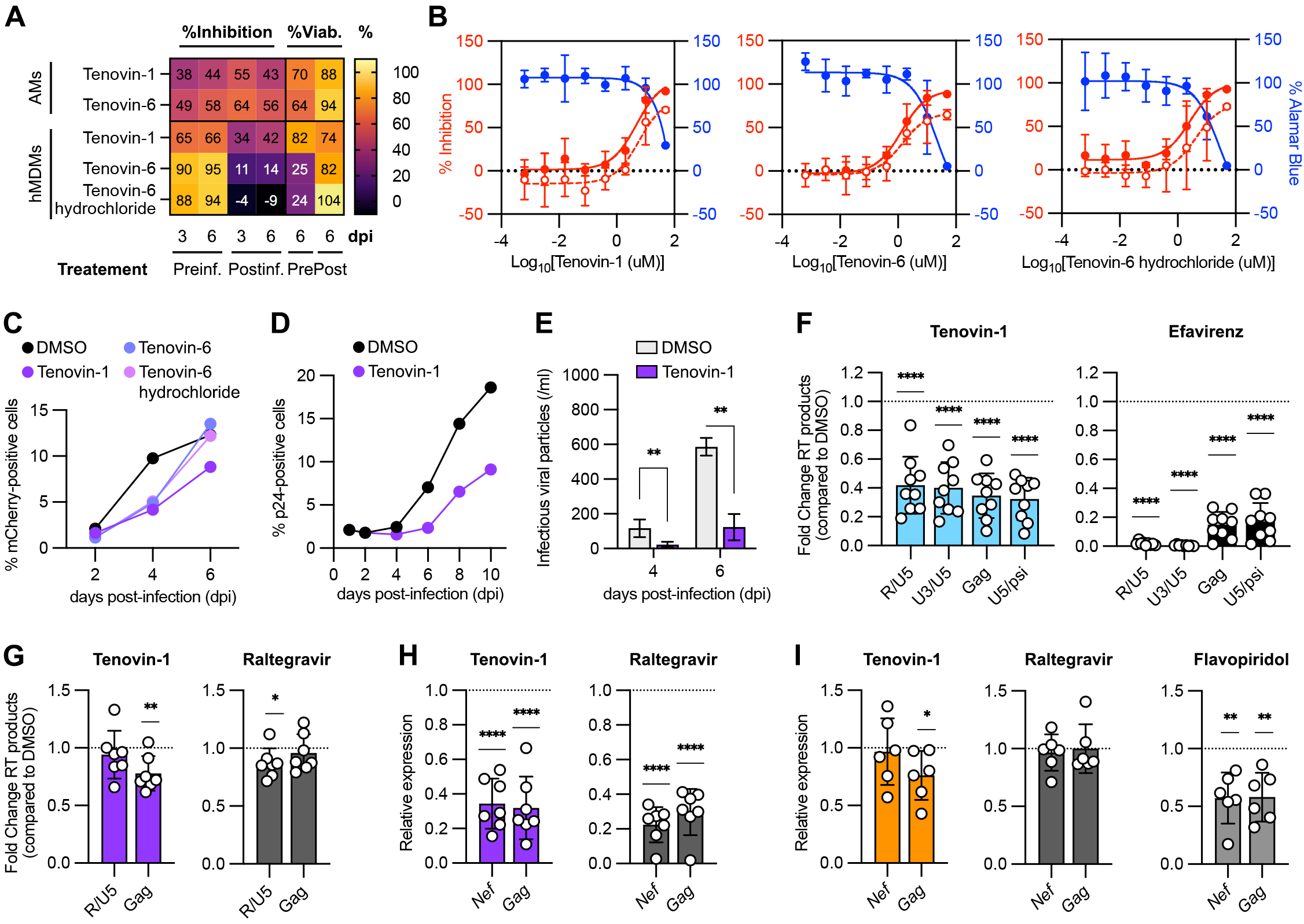
Tenovins block viral replication in human primary macrophages. (A-I) Macrophages were treated with Tenovin-1, -6 or -6 hydrochloride at 10mM one day before (A-B, F) or one day after (A, C-E, G-H), or 4 days after (I) HIV infection with NL4-3 gLuc + VSVg at MOI 0.1 (A-B) or MOI 0.2 (C-E); or NL4-3 Denv mCherry + VSVg at MOI 0.2 (F-I). (A) Gaussia luciferase was quantified at 3 and 6dpi, while Alamar blue was quantified at 6dpi, then percentage of inhibition and percentage of viability were plotted from 5 donors for hMDMs and from 3 donors for AMs. (B) Macrophages were treated with compounds at different concentration from 0.64nM to 50mM, then infected the next day. Percentage of inhibition at 3dpi (empty red circle) and 6dpi (full red circle) were plotted along with percentage of Alamar blue (full blue circle) for 2 donors. (C-D) At different time post-infection, macrophages were harvested, fixed, permeabilized and labeled with the antibody HIV-1 core antigen-RD1 KC57 (D), then analyzed by flow cytometry, for 1 donor. (E) At 4 and 6dpi, supernatants were collected to quantify infectious viral particles with TZM-bl assay, for 1 donor. (F) Macrophages were treated one day before HIV infection with Tenovin-1 (light blue) or Efavirenz at 5mM (black), as a positive control, then 2 dpi, DNA was extracted to quantify the different RT products (R/U5, U3/U5, Gag, U5/psi) by qPCR from 9 donors. (G-H) Macrophages were treated one day post-infection with Tenovin-1 (purple) or Raltegravir at 10mM (dark gray), as a positive control, then 2dpi, DNA and RNA were extracted from 7 donors to quantify RT products R/U5 and Gag by qPCR (G), and viral gene expression of *nef* and *gag* by RT-qPCR (H). (I) Macrophages were infected, then treated 4dpi with Tenovin-1 (orange), or Raltegravir at 10mM (dark grey) or Efavirenz at 5mM (black), as control. Six dpi, RNA was extracted from 6 donors to quantify viral gene expression of *nef* and *gag* by RT-qPCR.

In parallel flow cytometery assays using the non-replicative NL4-3 Denv mCherry reporter virus, we confirmed that Tenovin-1 is more effective at decreasing the number of HIV-positive cells than Tenovin-6 and Tenovin-6 hydrochloride when cells were treated at 1dpi (Fig 4C). To evaluate the efficacy of Tenovin-1 on viral spreading we infected hMDMs with HIV-gLuc pseudotyped with VSVg and treated with Tenovin-1 the following day (Fig 1C). Viral productivity was measured by quantifying both p24-positive cells and the production of infectious viral particles using flow cytometry and a TZM-bl reporter assay, respectively (Fig 4D-E). We observed that Tenovin-1 treatment inhibited viral replication up to 10 days post-infection (Fig 4D), leading to a marked decrease in production of infectious viral particles (Fig 4E).

We have also found that Tenovin-1 reduced reverse transcription products when hMDMs are treated one day before infection (Fig 3E-F). Upon further examination, we find Tenovin-1 reduced early products (R/U5) as well as all later RT products (U3/U5, Gag, U5/Y). These data suggest that Tenovin-1 inhibits viral reverse transcription at the step of initiation, as noted for EFV (Fig 4F). To assess the effect of Tenovin-1 and its activity after RT, hMDMs were infected with a single round virus (NL4-3 Denv mCherry) and treated with Tenovin-1 the next day (Fig 1C). Two days post infection and one day post treatment, RNA and DNA were extracted for quantification of viral RNA and RT products. We observed that Tenovin-1 slightly decreases RT products (Fig 4G), while it strongly inhibits the expression of the viral genes, *nef* and *gag* by 66% and 71%, respectively, levels comparable to Raltegravir treatment (Fig 4H). To evaluate the effect of Tenovin-1 on viral transcription, hMDMs were infected with the same single round virus, then treated at 4dpi, after integration event (Fig 2C). We observed that Tenovin-1 had minimal effect on *gag* expression (24% inhibition) and no effect on *nef* expression (Fig 4I), suggesting that Tenovin-1 does not inhibit viral transcription *per se* but appears to suppress steps between reverse transcription and viral transcription, which would potentially include integration.

In conclusion, the multitarget agent, Tenovin-1, shows lower cytotoxicity and more effective reduction of viral replication than its analogs (Tenovin-6 and Tenovin-6 hydrochloride), and appears to mediate its effect on inhibiting RT, and post-RT steps as opposed to impacting viral transcription.

### Host metabolic targets that regulate RT progression and post-RT steps of viral cycle in primary macrophages

Amongst the host cell inhibitory activities attributed to Tenovin-1 is its ability to inhibit dihydroorotate dehydrogenase (DHODH) (Ladds et al. 2020), a critical enzyme in *de novo* pyrimidine synthesis. To assess the possible MOAs of Tenovin-1 through DHODH activity, we selected several additional DHODH inhibitors: ML390, described as a potent inhibitor against enterovirus; Leflunomide and Brequinar, known to inhibit SARS-CoV-2 replication (Luganini et al. 2025). To identify the most potent compound inhibiting HIV replication in macrophages, we performed dose response experiments by pre-treating hMDMs with DHODH inhibitors prior to HIV-gLuc infection (Fig 3A). At 6dpi, Brequinar and ML390 showed an IC_50_ at 0.7mM and 7.4mM, respectively, while Leflunomide was less active (Fig 5A, Table S1). The data provide preliminary support for the hypothesis that DHODH activity is important for optimal viral replication in HIV-infected macrophages.

**Figure 5:**
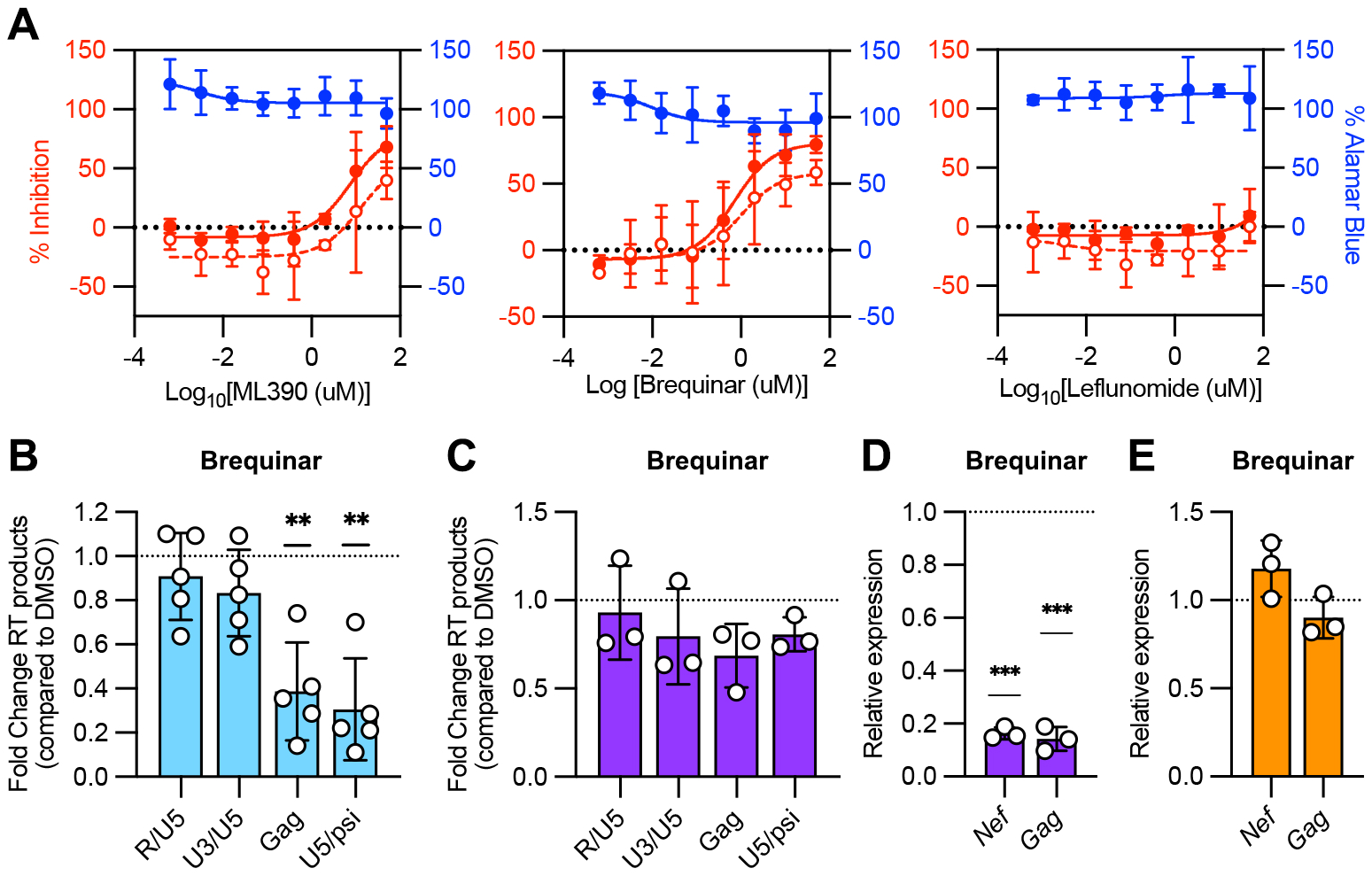
DHODH inhibition blocks viral replication in hMDMs. (A) Macrophages were treated with ML390, Brequinar or Leflunomide at different concentration from 0.64nM to 50mM, then infected the next day with NL4-3 gLuc + VSVg at MOI 0.1. Percentage of inhibition at 3dpi (empty red circle) and 6dpi (full red circle) were plotted along with percentage of Alamar blue (full blue circle) for 2 donors (ML390, Leflunomide) or 3 donors (Brequinar). (B-E) Macrophages were infected with NL4-3 Denv mCherry + VSVg at MOI 0.2, and treated one day before infection (B), one day after infection (C-D) or 4dpi (E) with Brequinar at 10mM. (B-C) Two dpi, DNA was extracted to quantify the different RT products (R/U5, U3/U5, Gag, U5/psi) by qPCR from 5 donors (B) or 3 donors (C). (D) Two dpi, RNA was extracted from 3 donors to quantify viral gene expression of *nef* and *gag* by RT-qPCR. (E) Macrophages were infected, then treated at 4dpi with Brequinar, then 6dpi, RNA was extracted from 3 donors to quantify viral gene expression of *nef* and *gag* by RT-qPCR.

We selected the most active compound, Brequinar, to further characterize the role of DHODH in HIV viral replication in macrophage. First, we analyzed RT efficacy with vDNA quantification by qPCR, after compound treatment and HIV infection with a non-replicative virus, using a protocol comparable to that detailed for Fig. 4F. In contrast to Tenovin-1, Brequinar did not reduce the amount of early RT products but did reduce the amount of later RT products (Fig 5B), indicating that DHODH activity is necessary for RT progression but not its initiation. When hMDMs were treated one day post-infection, and viral DNA and RNA quantified 2dpi similar to Fig 4G-H, the DHODH inhibitor exhibited minimal impact on RT products (Fig 5C). However, expression of *nef* and *gag* transcripts were strongly decreased by 83% (Fig 5D), suggesting DHODH activity is required in the post-RT steps of the viral cycle in macrophages. In contrast, when cells were infected with a non-replicative virus, then treated 4dpi (after integration) viral RNA expression was unchanged (Fig 5E), indicating that Brequinar does not inhibit viral transcription directly. These data suggest that DHODH activity is important for viral steps occurring between reverse transcription and viral transcription, such as integration, in primary macrophages.

In conclusion, in HIV-infected macrophages, DHODH activity is also of significance to RT progression, post-RT steps but not viral transcription.

### Inhibition of Sirtuin functions inhibits HIV-1 replication in primary macrophages

Another activity attributed to Tenovin-1 is its ability to activate p53 by protecting it from MDM2-mediated degradation through the inhibition of the protein deacetylating activity of sirtuins (SIRT1 and SIRT2) (Ladds et al. 2020; Lain et al. 2008). To assess these additional activities attributed to Tenovin-1, we selected further compounds from the original library: one SIRT1 inhibitor/p53 activator, Inauhzin ; one SIRT1/2 inhibitor, Salermide ; one SIRT1 inhibitor, Selisistat ; and one SIRT2 inhibitor, AK-7 (Table S1). We also selected an additional compound, Nutlin-3a, a specific p53 activator to test whether or not p53 could be implicated in the anti-HIV-1 activity (Breton et al. 2019). To determine both the cytotoxicity and inhibitory activity on viral productivity, we performed dose responses experiments by pre-treating hMDMs prior to HIV-gLuc infection (Fig 3A). The compounds Inauzhin, Selisistat and Salermide exhibited their strongest inhibition at 50mM with minimum cytotoxicity, while AK-7 and Nutlin-3a were more potent and less cytotoxic at 10mM (Fig 6A). These data indicate that both SIRT1/SIRT2 inhibition and p53 activation have the capacity to reduce viral replication in primary macrophages. However, the activities observed across the inhibitors tested suggest that inhibition may be multi-factorial and due to a combination of SIRT1/2 inhibition and/or p53 activation.

**Figure 6:**
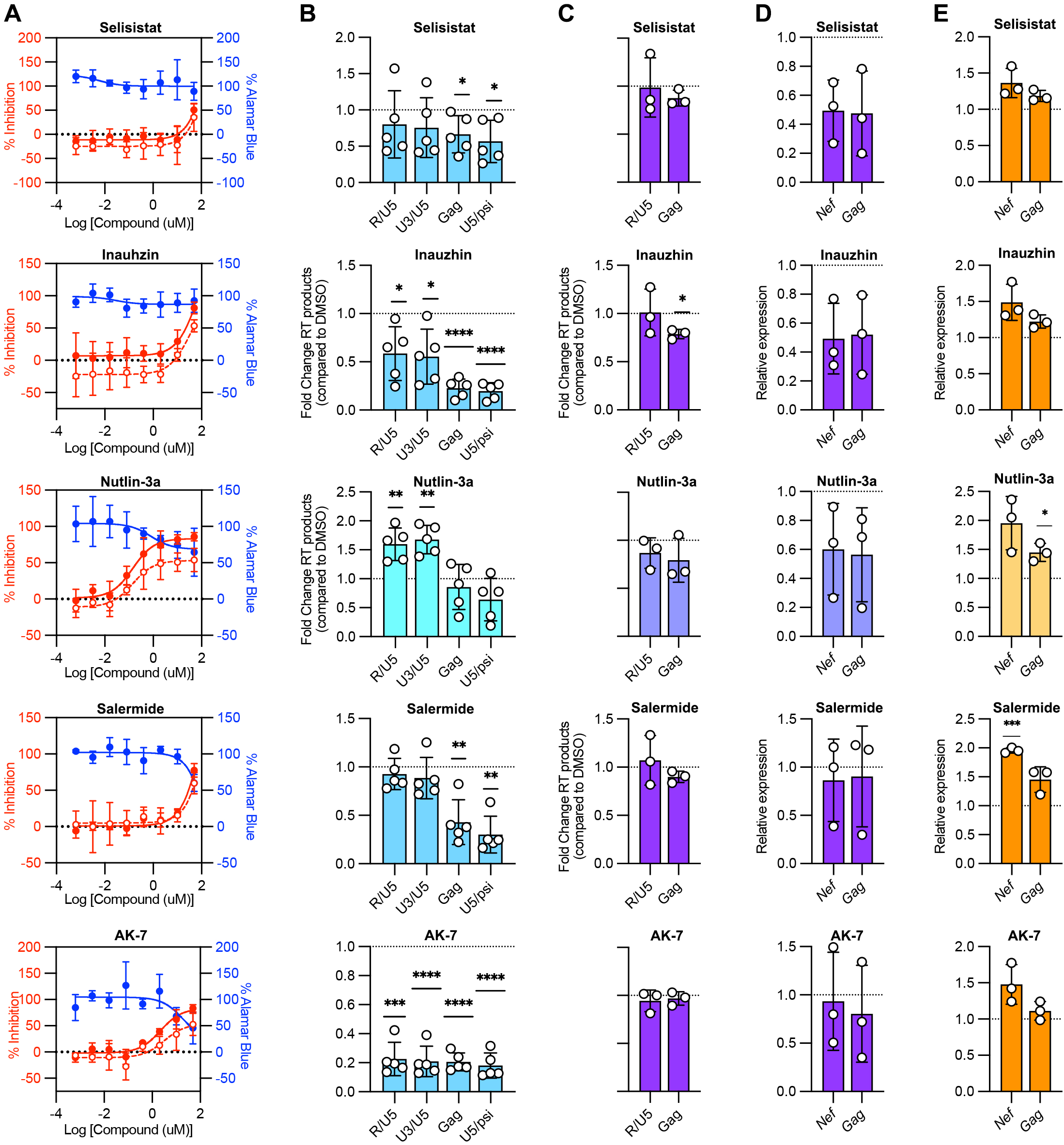
SIRT1/2 activity regulates reverse transcription in hMDMs. (A) Macrophages were treated with Selisistat (1^st^ row), Inauzhin (2^nd^ row), Nutlin-3a (3^rd^ row), Salermide (4^th^ row) or AK- 7 (5^th^ row) at different concentration from 0.64nM to 50mM, then infected the next day with NL4- 3 gLuc + VSVg at MOI 0.1. Percentage of inhibition at 3dpi (empty red circle) and 6dpi (full red circle) were plotted along with percentage of Alamar blue (full blue circle) for 3 donors. (B-E) Macrophages were infected with NL4-3 Denv mCherry + VSVg at MOI 0.2, and treated one day before infection (B), one day after infection (C-D) or 4dpi (E) with Selisistat at 50mM (1^st^ row), Inauzhin at 50mM (2^nd^ row), Nutlin-3a at 10mM (3^rd^ row), Salermide at 50mM (4^th^ row) or AK-7 at 10mM (5^th^ row). (B-C) Two dpi, DNA was extracted to quantify the different RT products (R/U5, U3/U5, Gag, U5/psi) by qPCR from 5 donors (B) or 3 donors (C). (D) Two dpi, RNA was extracted from 3 donors to quantify viral gene expression of *nef* and *gag* by RT-qPCR. (E) Macrophages were infected, then treated at 4dpi with compounds, then 6dpi, RNA was extracted from 3 donors to quantify viral gene expression of *nef* and *gag* by RT-qPCR.

To attempt to discriminate between the cellular targets involved in the inhibition of RT activity observed following Tenovin-1 treatment, hMDMs were pretreated with the compounds reported to target p53, SIRT1 and SIRT2 independently, then infected with a non-replicative virus. Two days later viral DNA was quantified, using a timeline comparable to that of Fig 4F and Fig 5B. Similar to the Tenovin-1 effect on RT, the SIRT2 inhibitor AK-7 strongly reduced all RT products from early (R/U5) to late (U5/Y) intermediates (Fig 6B), revealing the involvement of SIRT2 activity in the initiation of reverse transcription. In contrast, the inhibitors of SIRT1 (Selisistat, Salermide, Inauzhin) reduced the late RT products Gag and U5/Y (Fig 6B), suggesting that these compounds impacted the progression of reverse transcription, rather than its initiation. Surprisingly, p53 activation induced by Nutlin-3a lead to an increase of early RT products R/U5 and U3/U5 (Fig 6B). These data suggest that SIRT2 activity is important for RT initiation while SIRT1 is necessary for its progression.

To determine which activity of Tenovin-1 is responsible for inhibiting the post-RT steps observed in Fig 4G-H, hMDMs were treated with the different compounds 1dpi, then viral DNA and viral RNA were quantified 2dpi, as shown in Fig 4G-H and Fig 5C-D. None of these compounds reduced the amount of RT products (Fig 6C), and only the SIRT1 inhibitors and p53 activators, Selisistat, Inauzhin and Nutlin-3a, led to a decrease of viral productivity (Fig 6D), suggesting a role for SIRT1 and p53 in post-RT steps of the viral cycle. None of these compounds reduced viral productivity when cells were treated at 4dpi, after integration (Fig 6E). These data imply that SIRT1 and p53 mediate their activity in primary macrophages on viral steps occurring between reverse transcription and viral transcription, which would include integration.

These results indicate that Tenovin-1 has pleotropic inhibitory activities on different stages in the viral life cycle. The compound inhibits viral cycle steps between RT and viral transcription potentially through p53 activation and/or SIRT1 and DHODH inhibition, and the inhibition of DHODH and SIRT activities may also reduce reverse transcription.

## DISCUSSION

Since the introduction of antiretroviral therapy (ART), a combination of antiretroviral drugs targeting HIV proteins, the life expectancy of HIV-infected patients has significantly increased. However, using these direct-acting agents (DAA) can lead to resistance, treatment failure, or viral rebound if the treatment is discontinued (Chun et al. 2013). Despite the impressive success of drug development programs there are two specific areas in which the progress has been challenged. Firstly, there is no obvious “Road to Cure” because all current therapies suppress new infections without resolving pre-existing infections. Secondly, most ongoing anti-HIV-1 drug programs are interrogating known targets and while these programs will undoubtedly lead to better drugs, they still fall short of addressing the central issues of persistent infection (Pierson, McArthur, and Siliciano 2000). We believe this represents a downside of moving from biological screening, or discovery, to target based drug development, which excludes the broader biology of viral infection. Amongst the cell types responsible of HIV persistence, macrophages have tended to be understudied. However, over the past decade, accumulating evidence have highlighted significant participation of macrophages in viral transmission and dissemination, with tissue-resident macrophages emerging as sites of viral persistence in people living with HIV-1 (Hendricks et al. 2021; Veenhuis et al. 2021). These cells are susceptible to HIV-1 infection, and appear capable of sustaining viral genomes in ART-suppressed individuals, however, they have received little attention in the area of drug discovery. The primary goal of the current study is to identify cellular pathways that had not previously been implicated in HIV-1 productivity in primary macrophages. We believe this represents a viable route to the identification of novel target candidates for further characterization and evaluation. While the scope of our screen is limited in its chemical diversity, we have deliberately focused compounds implicated in epigenetic control of the host cell, including some compounds known to modulate HIV latency (Jones et al. 2026).

The recent therapeutic approaches involving host-targeting agents (HTA) have been extensively studied to target latently infected cells by identifying latency reversing agents (LRA) or latency promoting agents (LPA) for “shock-and-kill” and “lock-and-block” strategies, respectively (Tanaka et al. 2022; Vansant et al. 2020). Most of these screens have been performed in T cells (Jones et al. 2007; Yang et al. 2009; Tioka et al. 2025), however, a recent screen identified compounds that control HIV silencing in hMDMs using a single round infection (Yi et al. 2023). With this in mind we designed our screen differently to address two aspects of HIV-1 infection not included in early studies. First, our screen design involves a spreading viral infection with intact virus, meaning that the entire viral life cycle is interrogated, not only the viral transcription. Second, our screen is conducted on two primary human macrophage models: monocytes-derived macrophages differentiated *in vitro*, and tissue resident alveolar macrophages from the human lung, known to potentially play a crucial role in HIV persistence (Costiniuk et al. 2024).

Our macrophage-based screen identified 21 verified HTAs that controlled different steps of the viral cycle in primary macrophages, including inhibitors/activators of kinases (AMPK, PKC, AURK, JAK) and transcription factors (HIF, p53); inhibitors of methyl transferases (DNMT, HMT), demethylases (HDM), acetyl transferases (HAT), deacetylases (SIRT), polymerase (PARP) and bromodomain (BRD) (Fig 7). Some of the compounds have already been identify as inhibitors of viral latency and viral transcription (Jones et al. 2026), which we view as further validation of both studies. Among them, the PKC agonist PDBu was reported to reactivate latent viral infection in T cells while suppressing the viral replication in monocytic cell lines (Kalvatchev, Walder, and Garzaro 1997). Decitabine, a DNMT inhibitor, has been reported to reactivate HIV-1 gene expression when used in combination with LRAs, through modulation of chromatin accessibility, and to also exhibit antiretroviral activity against HIV-1 through lethal mutagenesis (Blazkova et al. 2009; Kauder et al. 2009; Bouchat et al. 2016; Clouser et al. 2014; Clouser, Chauhan, et al. 2012; Clouser, Holtz, et al. 2012; Clouser, Patterson, and Mansky 2010; Rawson, Daly, et al. 2016; Rawson, Roth, et al. 2016). The BRD4 inhibitor JQ1, and parent molecule of dBET1 and PROTAC BET Degrader-10 (PBD-10), has been described to activate HIV transcription and reverse latency in various cell types, including macrophages (Zhu et al. 2012; Banerjee et al. 2012; Li et al. 2013; Boehm et al. 2013; Kisaka et al. 2024). The PARP-1 inhibitor PJ34 has been shown to induce a reduction in LTR activity in macrophages (Rom et al. 2015), while CBP/p300 (HAT) inhibitors, GNE-049 and CPI-637 have been described as LRAs (Zheng et al. 2021; Lindqvist et al. 2020). As in these previous studies, we have shown that some of these compounds regulate viral transcription in human primary macrophages, but our data also indicate inhibition of reverse transcription, suggesting a dual mechanism of action and a strong potential for therapy (Figure 7, Table S1).

**Figure 7:**
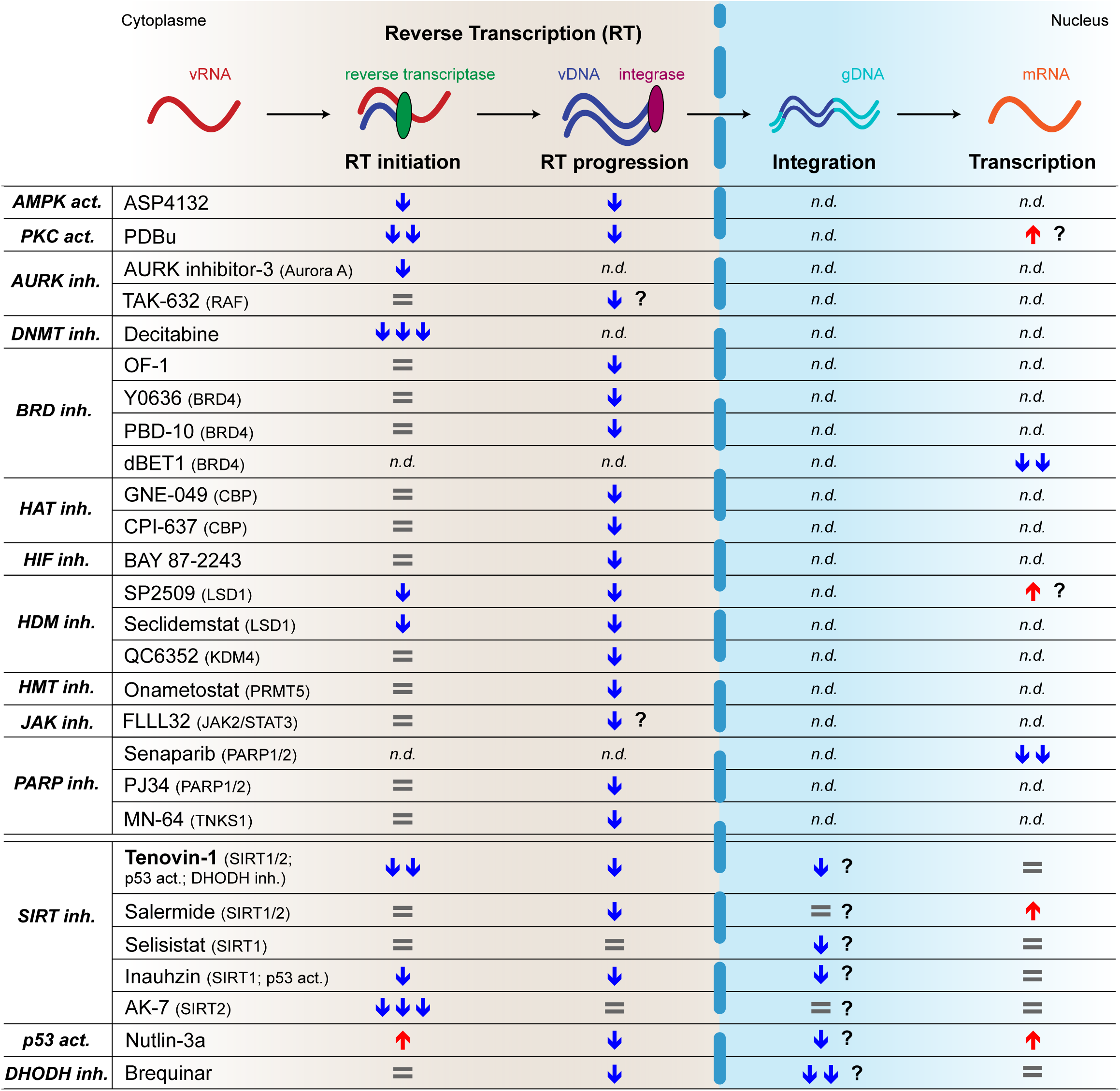
Summarize of compounds regulating different steps of viral cycle in HIV-infected hMDMs.

Most of the epigenetic targets emerging this screen have been described as central regulators in viral transcription and latency (Jones et al. 2026), including AMPK, Raf/AURK, LSD1, SIRT1/2 and p53 in various cell types (Zhang et al. 2012; Vargas et al. 2019; Sakane et al. 2011; Le Douce et al. 2012; Pagans et al. 2005; Zhang and Wu 2009; Zhang et al. 2009; Samer et al. 2020; Duran-Castells et al. 2023; Mukerjee et al. 2010). This study demonstrates that these epigenetic targets play a role in the HIV life cycle within macrophages. Indeed, we identified several compounds inhibiting viral reverse transcription (Fig 3E-F, Fig 7), such as ASP4132 (AMPK activator); AURK inhibitor-3 and TAK-632 (Raf/AURK inhibitors); SP2509 and Seclidemstat (LSD1 inhibitors); Tenovin-1 (SIRT1/2 inhibitor, p53 activator). Notably, this last compound, which has also been reported to inhibit DHODH (Ladds et al. 2020), exhibits broad-spectrum antiviral activity against arboviruses including flaviviruses and bunyaviruses (Wan et al. 2021; Hackett et al. 2019; Rausch et al. 2017), a list of viruses to which HIV may now be added.

The compounds identified in our infected macrophage screen represent a valuable resource for future investigations, providing new insights into potential points of interference at multiple stages of the HIV replication cycle, beyond viral transcription, in human primary macrophages. Among them, some FDA-approved drugs, such as Decitabine and Senaparib, could be explored for potential repurposing against HIV (Kim et al. 2022; Wu et al. 2024). Collectively, our data identify a broad range of host-acting agents with differing modes of antiviral activities that we believe provides a valuable foundation for the development of novel therapeutic strategies against HIV.

## MATERIAL & METHODS

### Primary human macrophages

Human monocyte-derived macrophages (hMDMs) were obtained from human monocytes from peripheral blood mononuclear cells of healthy individuals by counter current centrifugal elutriation with an average purity of >97% (Elutriation Core Facility, University of Nebraska Medical Center). For differentiation in macrophages, monocytes were maintained in DMEM media (Corning, #15- 017-CV) supplemented with 10% human serum (LGC Diagnostics/Seracare, #1830-0003), 2mM L-glutamine (Corning, #25-005-CI), 10mM HEPES (Fisher Scientific, #15630080), 1mM Sodium Pyruvate (Corning, #25-000-CI), 100IU/mL penicillin and 100μg/mL streptomycin (Corning, #30- 002-CI), for 6 days at 37°C with 6% CO_2_.

Human alveolar macrophages (AMs) were obtained from bronchoalveolar lavage (BAL) from healthy participants individuals (aged ≥18 y) at the Queen Elizabeth Central Hospital in Blantyre, Malawi. The study received ethical approval from the research ethics committees of the Malawi National Health Sciences Research Committee (Research protocol Number #22/04/2904 ), the Liverpool School of Tropical Medicine, UK (Research protocol 08.54), and Cornell University (Research protocol 908000698). All participants provided written informed consent. AMs were maintained in RPMI 1640 media with L-glutamine (Gibco, #3256089) supplemented with 10% Fetal Bovine Serum (Sigma Aldrich, #0001669683), 10mM HEPES(Gibco, #2744493), Penicillin-Streptomycin 10,000U/mL (Gibco, #225194).

### Virus production

The plasmids encoding HIV-1 molecular clones NL43-IRES-Gaussia Luciferase with R5-tropic env (HIV-gLuc) and NL43-IRES-mCherry Denv were generated by replacing the mCherry sequence with that of Gaussia Luciferase or by removing *env* sequence from original plasmid pNL43-IRES-mCherry R5-env (BaL) used in (Boliar et al. 2019), respectively. HIV-1 viruses pseudotyped with VSV-G were prepared by cotransfecting HEK-293FT cells (ATCC, #CRL-3216) with one of the HIV-1 molecular clones and VSV-G expression plasmid using Lipofectamine 3000 reagent (Life Technologies, #L3000015) in Opti-MEM media (Life Technologies, Gibco #31985070). The next day, transfection media was replaced with fresh antibiotic free DMEM media supplemented with 10% FBS (Harvest Midsci USFBS), 2mM L-glutamine, 10mM HEPES and 1mM Sodium Pyruvate, the next day. Then 48h posttransfection, cell culture supernatant containing HIV-1 virus was harvested, centrifuged to remove cell debris, filtered using 0.45-μm filter, aliquoted and stored at −80 °C. Viral titers were determined by infection of the indicator cells HeLa TZM-bl (bearing the β-galactosidase gene under the control of HIV-1 LTR; from NIH Reagent Program) with serial dilutions of the stocks, followed by a β-galactosidase coloration of the cells and counting of blue cells.

### Macrophage-based screening

The epigenetic compounds library from MedChemExpress company included 721 small molecules diluted at 2mM in DMSO (MedChemExpress, #HY-LD-000002646), and Efavirenz, Raltegravir, Flavopiridol were used as positive control. Macrophages were plated at 500,000 cells/well in white, optical-bottom tissue culture-treated 96-well plates (Nunc^TM^, #12-566-71), treated with epigenetic compounds at 10mm one day before infection (Figure 3), one day after infection (Figure 1) or 4 days after infection (Figure 2). Macrophage were infected using HIV-gLuc pseudotyped with VSVg at MOI 0.1 for 24h, the next day the media was replaced with fresh hMDMs media +/- epigenetic compounds. Six days post-infection, 10ul of supernatant was collected and diluted 10 times into Nunc^TM^ non-treated white-bottom 96-well plates (Nunc^TM^, #12- 566-02) to measure Gaussia Luciferase activity using Pierce™ Gaussia Luciferase Glow Assay Kit (Thermo Scientific, #PI16161) and by following the manufacturer’s instructions. In parallel, 10ul of Alamar Blue HS Cell Viability Reagent (Invitrogen, #A50101) was added on the cells, and after 60min, fluorescence (570/590nm) was measured using the Envision plate reader (PerkinElmer), also used for luminescence reading. Percent of inhibition (luciferase expression) was calculated from Relative luminescence unit (RLU) values where 0% was equal to the RLU values in the presence of DMSO, and 100% was equal to the RLU from uninfected wells (see below).

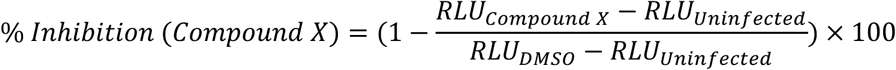

Percent of Alamar blue was calculated from Relative fluorescent unit (RFU) values where 100% was equal to the RFU in the presence of DMSO only (see below).

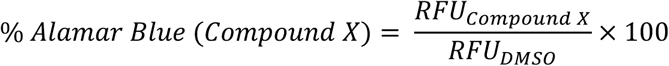

### Flow cytometry

To assess viral infection of hMDMs, cells were harvest at different time point post-infection (HIV- mCherry Denv+VSVg or HIV-gLuc+VSVg at MOI 0.2), by gentle scraping after incubation in cold PBS and then fixed in 1% paraformaldehyde in PBS. For detection of HIV-infected hMDMs, cells were permeabilized with 0.05% Saponin then stained with an antibody against the HIV-1 core protein p24 (Beckman Coulter, KC57-RD1) for 30 min at room, or mCherry-positive cells were directly quantified with Attune Nxt Flow cytometer (ThermoFisher). Data were analyzed using FlowJo software with the following gating strategy: cells were first gated for macrophage (side scatter area - SSC-A *vs* forward scatter height area - FSC-A), then singlets (FSC-H *vs* FSC-A) and the frequency of HIV-positive hMDMs was calculated based on HIV-1 p24-PE or mCherry expression.

### qPCR and RT-qPCR

For viral expression analysis, hMDMs were infected with the single-round VSV-G pseudotyped NL43–IRES-mCherry-Denv virus (MOI 0.2) and treated with compounds at 1dpi or 4dpi. At 2 or 6dpi, total RNA was extracted using TRIzol reagent (Invitrogen, #15596026) or AllPrep DNA/RNA kit (Qiagen, #80204) according to manufacturer’s instructions, followed by DNase treatment using Turbo DNA-free kit (Invitrogen, #AM1907) to remove genomic DNA. The same amount of RNA was used for cDNA synthesis and subsequent quantification of viral expression using ABI 7500 Fast Real-time PCR system (Applied Bioscience) following manufacturer’s protocol. HIV-1 viral transcripts were quantified by RT-qPCR using iScript cDNA Synthesis kit (Bio-Rad, #1708890) and iTaq Universal SYBR Green Supermix (Bio-Rad, #1725120) for Gag (Forward primer: AAG CAC TGG GAC CAG GAG C and Reverse primer: TGG TAG CTG GAT TTG TTA CTT GGC) and Nef (Forward primer: TAG TGT GAT TGG ATG GCC TGC and Reverse primer: ACA AGC ATT GTT AGC TGC TG) according to manufacturer’s protocol. Expression of the housekeeping genes was used for normalization of RT-qPCR expression data: GAPDH (Forward primer: GAC AAG CTT CCC GTT CTC AG and Reverse primer: GAG TCA ACG GAT TTG GTC GT), U6 (Forward primer: CTC GCT TTG GCA CA and Reverse primer: AAC GCT TCA CGA ATT TGC GT), and 18S rRNA (Forward primer: GGC CCT GTA ATT GGA ATG AGT C and Reverse primer: CCA AGA TCC AAC TAC GAG CTT).

For analysis of reverse transcription products (RT), hMDMs were infected with the single-round VSV-G pseudotyped NL43–IRES-mCherry-Denv virus (MOI 0.2) and treated with compounds at 1 day prior or 1 day post infection (dpi). At 2dpi, total DNA was extracted using DNeasy Blood & Tissue Kit (Qiagen, #69504) or AllPrep DNA/RNA kit (Qiagen, #80204) according to manufacturer’s instructions. The same amount of DNA was used for vDNA quantification using ABI 7500 Fast Real-time PCR system (Applied Bioscience) following manufacturer’s protocol. HIV-1 viral RT were quantified by qPCR using iTaq Universal SYBR Green Supermix (Bio-Rad, #1725120) for Early RT (Forward primer: GTG CCC GTC TGT TGT GTG AC and Reverse primer: GGC GCC ACT GCT AGA GAT TT), Jump strand transfer or U3/U5 (Forward primer: GTG CCC GTC TGT TGT GTG AC and Reverse primer: GGC GCC ACT GCT AGA GAT TT), RT intermediate or Gag (Forward primer: GAG CCC TCA GAT GCT GCA TAT and Reverse primer: CCA CAC TGA CTA AAA GGG TCT GAG), Late RT or U5/psi (Forward primer: TGT GTG CCC GTC TGT TGT GT and Reverse primer: GAG TCC TGC GTC GAG AGA GC) based on (Mbisa et al. 2009) and according to manufacturer’s protocol. Housekeeping gene GAPDH was used for normalization of qPCR data (Forward primer: GAC AAG CTT CCC GTT CTC AG and Reverse primer: GAG TCA ACG GAT TTG GTC GT).

### Statistical analysis

Statistical analysis was performed using GraphPad Prism software, one-way ANOVA with Tukey’s multiple comparison test was used for all statistical analysis with a *p*-value below 0.05 considered as significant.

## ACKNOWLEDGMENTS

This work was supported by NIH grant AI176575 to DGR. This portion of the work performed in Malawi was supported through a sub-contract in AI176575 that was terminated in 2025.

