## Supplementary figures and images for "Screening of epigenetic modifiers identifies novel host-directed agents that suppress HIV-1 replication in primary human macrophages"

### Supplementary Figures

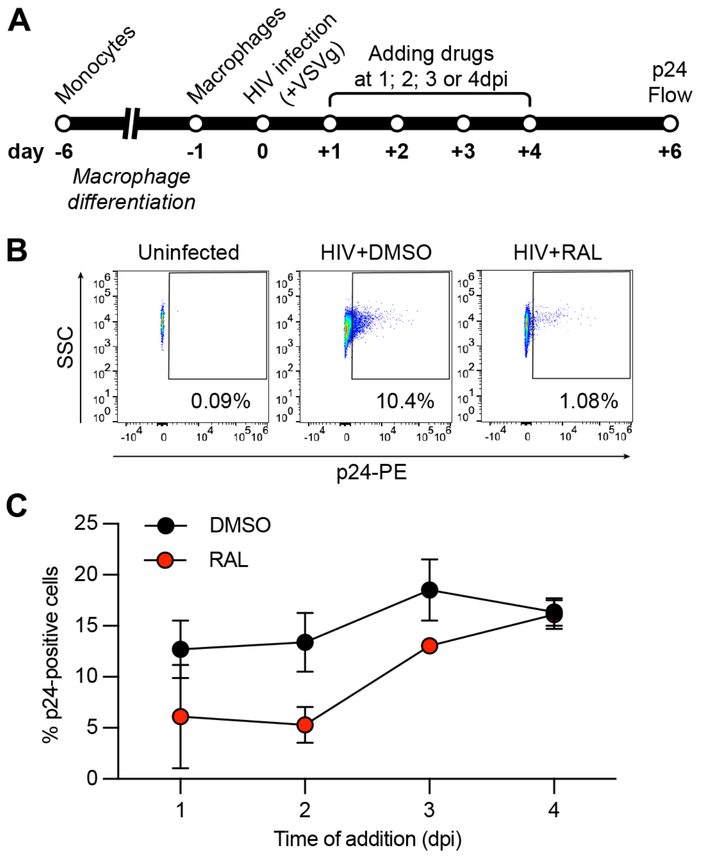


**Supp Figure 1**

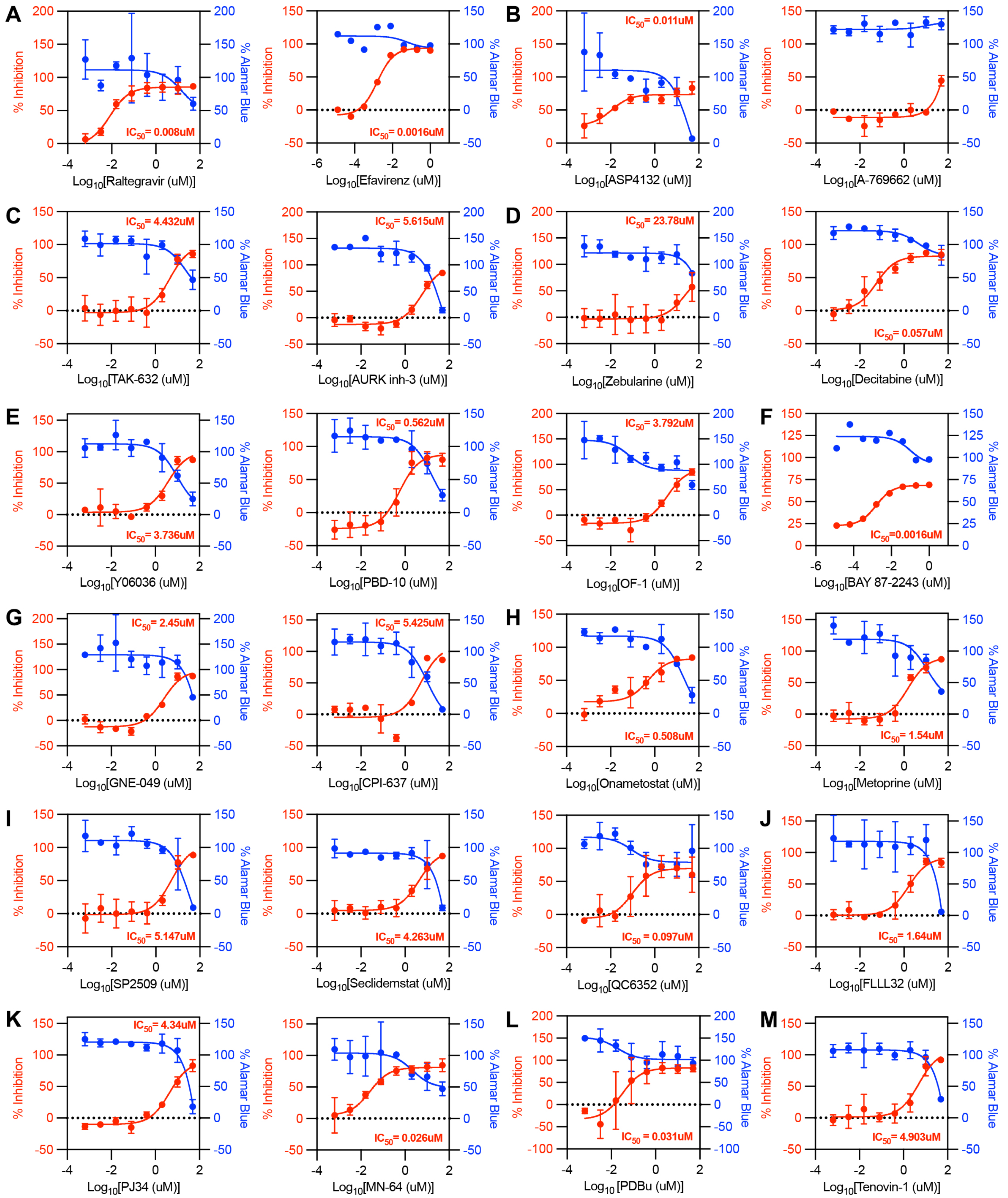


**Supp Figure 2**
